# Repeated Sub-anaesthetic dose of Ketamine Elevates Superoxide Dismutase in Pharmacological Model of Schizophrenia-Like Phenotypes in Mice

**DOI:** 10.1101/2024.08.28.610179

**Authors:** Usman Yusuf, Aderibigbe Oladele Adegbuyi, Fehintola Fatai Adewale

## Abstract

**Objective:** The study evaluated behavioural phenotypes and the level of Superioxde Dismutase activity in repeated sub-anaesthetic dose of ketamine administered to model schizophrenia in animal study.

**Method:** Animals were divided into groups (n=6), control group received distilled water (10mL/kg) as vehicle (VEH), Ketamine treated group (KET) received sub-anaesthetic dose of ketamine (20mg/kg) for 14 days consecutively. Animals in risperidone treated group (KET+RISP) were pre-treated with sub-anaesthetic dose of ketamine (20mg/kg) alone for 7 consecutive days, and from day 8-14, risperidone (0.5mg/kg) was administered 1 hour post-ketamine treatment. All treatment were administered intraperitoneally (i.p). Twenty-four (24) hours after the last treatment, Behavioural phenotypes (locomotor activity and cognition) were assessed in locomotor activity cage and elevated maze plus. Thereafter level of superioxde dismutase (SOD) activity was evaluated in homogenized brain tissue of each mouse using spectrophotometric analysis.

**Result:** KET group showed significant (p<0.05) increase in movement counts and number of rearing events in locomotor activity test, also prolonged latency to enter the open arms in cognitive assessment compared to animals that received distilled water (10mg/kg), and risperidone (0.5mg/kg) treatment. The level of Superioxide Dismutase (SOD) activity was significantly elevated in KET group compare with vehicle control and risperidone treated animals.

**Conclusion:** Repeated dose of ketamine may pose differential effect on endogenous antioxidant system which may elevate superioxide dismutase activity in ketamine induced schizophrenic rodent as positive control mechanism.

## Introduction

Superoxide dismutase (SOD) is a key antioxidant enzyme that plays a critical role in dismutating superoxide radicals into hydrogen peroxide and oxygen, thus protecting cells from oxidative damage [1]. In schizophrenia, altered SOD activity has been observed, with studies showing both increased and decreased levels, depending on the stage of the disease and treatment status, with increased levels in first episode of schizophrenia patients [2]. The changes of antioxidant enzymes are related to schizophrenia and are part of its pathophysiology [3]. Padurariu et al. reported increased superoxide dismutase (SOD) activity in patients with schizophrenia [4], showing SOD activity is positively associated with both positive and negative symptoms, as well as general psychopathology in chronic schizophrenia patients [2].

Ketamine, an NMDA receptor antagonist, is commonly used in preclinical research to model schizophrenia in animals. Its sub-anesthetic doses have been shown to induce a variety of schizophrenia-like symptoms, including cognitive deficits, social withdrawal, and behavioural changes, which mimic the positive, negative, and cognitive symptoms of the disorder in humans [5]. Acute or repeated administration of ketamine is widely used animal model of schizophrenia [6, 7]. The drug’s action on glutamatergic neurotransmission, specifically its antagonism of NMDA receptors, leads to an increase in dopamine release, which is thought to underlie some of these effects [8].

Chronic exposure to sub-anesthetic dose of ketamine has been shown to elevate oxidative stress markers, including increased SOD activity due to activation in neurons of reduced nicotinamides adenine dinucleotide phosphate (NADPH) oxidase [9], also as compensatory response to oxidative damage [2]. This suggests that ketamine-induced schizophrenia models not only replicate the behavioural symptoms of the disorder but also the underlying biochemical abnormalities, such as altered endogenous anti-oxidant system. In animal model of schizophrenia, risperidone has been found to reduce the level of SOD [10]. Neuroprotective properties exhibited by risperidone could be attributed to microglial activation or antioxidant maintenance mechanisms in the cerebral cortex and hippocampus [11]. The study evaluated ketamine induced behavioural phenotypes and altered Superoxide Dismutase activity level in mice.

## Materials and Methods

### Animals

Male and female mice, aged 8-10 weeks, were used in this study. The animals were housed under standard laboratory conditions with a 12-hour light/dark cycle and ad libitum access to food and water. All experimental procedures were approved by the Institutional Animal Care and Use Committee (IACUC) and adhered to the guidelines set forth by the National Institutes of Health (NIH) for the care and use of laboratory animals.

### Drugs and Chemicals

Ketamine hydrochloride (Ranbaxy Pharm), Risperidone, carbonate buffer (pH 10.2), phosphate buffer (0.1 M, PH 7.4), adrenaline.

### Experimental Design

The mice were randomly assigned to control group, ketamine group or treatment group. The ketamine group received intraperitoneal injections of ketamine hydrochloride at a sub-anesthetic dose of 20 mg/kg once daily for 14 consecutive days. The control group received an equivalent volume of saline. The treatment group were pre-treated with ketamine alone (20mg/kg i.p) from day 1 to 7, and from the 8^th^ to the 14^th^ day received risperidone (0.5mg/kg i.p) treatment, 1 hour post-ketamine administration. The dose and regimen are based on previous studies demonstrating the induction of schizophrenia-like behaviours and altered oxidative stress [12].

### Behavioural Assessments

Twenty-four (24) hours after the last treatment, each mouse was subjected to behavioural tests to assess schizophrenia-like symptoms, including the open field test for hyperlocomotion, and numbers of rearing, while cognitive deficits was evaluated using elevated maze plus test, as previously described respectively with some modifications [13-15]

### Biochemical Analysis

Immediately after the behavioural tests, each mouse was sacrificed by cervical dislocation and decapitated, the brain tissues were removed, Thereafter, the whole brain tissues of each mouse was homogenized with 5 mL of 10% w/v phosphate buffer (0.1 M, PH 7.4). Brain tissue homogenates were centrifuged at 10,000 g for 10 min at 4°C, the pellet discarded and the supernatant portions were used for superoxide dismutase activity assay. The level of SOD activity was measured by the method described by Misra and Fridovich [12]. This method is based on the inhibition of superoxide dependent adrenaline auto-oxidation in a spectrophotometer adjusted at 480 nanometer (nm). A unit of SOD activity was given as the amount of SOD necessary to cause 50% inhibition of the oxidation of adrenaline and express as mmol/min/mg of tissue [12].

## Statistical Analysis

Data were expressed as Mean ± S.E.M and were analyzed using one–way analysis of variance (ANOVA) and post hoc tests (Student’s Newman-Keuls) for the multiple comparisons where appropriate using GraphPad InStat® Biostatistics software. The level of significant was set at p< 0.05

## Results

Locomotor activity assessed by measuring movement counts/5 minutes (in seconds), and the number of rearing events showed KET group exhibited a significant (p<0.05) increase in movement counts/5 minutes and number of rearing in open field test compare to VEH and KET + RISP mice. Specifically, KET showed an average movement of 549.0 ± 19.9, and 16.40 ± 0.93 rearing. The control showed an average movement of 377.4 ± 14.6, and 9 ± 0.71 rearing while the risperidone treated animals showed average movement of 446.0 ± 10.2 and 10.40 ± 0.93 rearing respectively (Tab. 1). Regarding cognitive function, the KET group showed a significant (p<0.05) increase in latency to enter the open arms with mean value of 34.30±1.77 seconds, in contrast to 18.79±0.47 and 24.36± 1.75 seconds in the control and risperidone treatment groups respectively (Tab. 2). KET mice significantly (p<0.05) showed elevated SOD activity, compared to vehicle control (VEH) and risperidone treated (KET + RISP) mice (fig.1).

**Table 1.** Locomotor Activity Assessments. Ketamine induced hyperlocomotion Values represents Mean ± Standard Error of Mean (n=5). One way ANOVA, significant [*F* (2.00, 9.552) = 31.43 *P*< 0.0001, *F* (2.000, 11.40) = 20.88, *P*<0.0001]. ^**^*p*< 0.05 as compared with KET group. **VEH**=Vehicle, **KET**=Ketamine, **RISP**=Risperidone

| S/N | Treatments | Movement counts/5 minutes in Activity cage (secs) | Numbers of Rearing |
| --- | --- | --- | --- |
| 1. | VEH | 377.4 ± 14.6<sup>**</sup> | 9 ± 0.71<sup>**</sup> |
| 2. | KET | 549.0 ± 19.9 | 16.40 ± 0.93 |
| 3. | KET + RISP | 446.0 ± 10.2<sup>**</sup> | 10.40 ± 0.93<sup>**</sup> |

**Table 2.** Evaluation of cognitive function. Ketamine induced Cognitive deficit.Values represents Mean ± Standard Error of Mean (n=5). One way ANOVA, significant [*F* (2.000, 8.551) =28.95, *P*<0.0001] *P*< 0.0001]. ^**^*p*< 0.05 as compared with KET group. **VEH**=Vehicle, **KET**=Ketamine, **RISP**=Risperidone

| S/N | Treatments | Latency period to enter open arms (5 minutes) in EMP (secs) |
| --- | --- | --- |
| 1 | VEH | 18.79±0.47<sup>**</sup> |
| 2 | KET | 34.30±1.77 |
| 3 | KET + RISP | 24.36± 1.75<sup>**</sup> |

**Figure 1.**
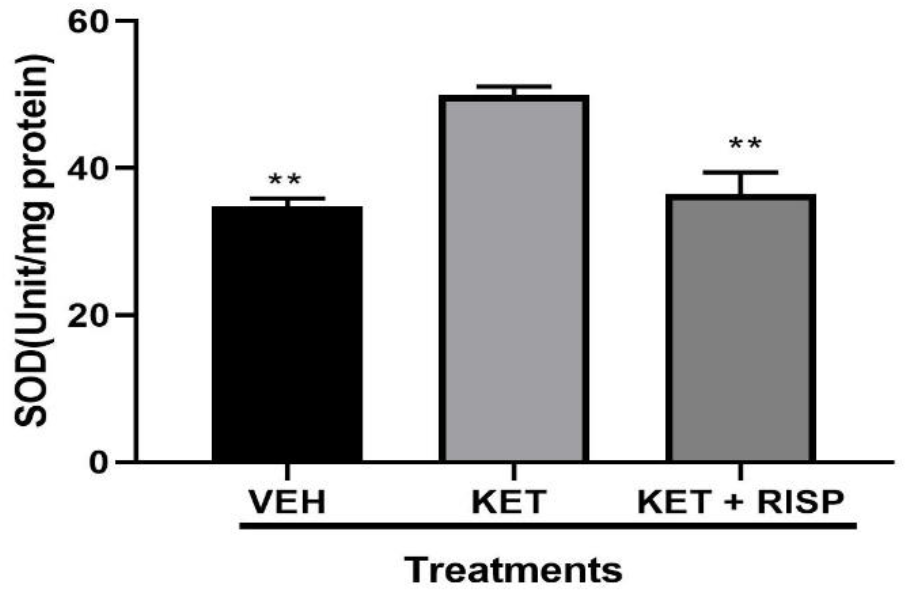
Evaluation of Superoxide Dismutase Activity Level. Superioxide Dismutase activity in homogenized mice brain. Values represents Mean ± Standard Error of Mean (n=5). One way ANOVA, significant [F (2.000, 9.055) =67.55, *P*< 0.0001]. ^**^*p*< 0.05 as compared with KET group. **VEH**=Vehicle, **KET**=Ketamine, **RISP**=Risperidone

## Discussion

Animal models, particularly rodent models, are widely used to study the biochemical and behavioural aspects of schizophrenia. In these models, alterations in brain biochemical activities, including compensatory enzymatic activities of superoxide dismutase, have been noted following the administration of psychotomimetic drugs like ketamine [16], coupled with schizophrenia-like behavioural manifestations [6]. In the context of this study, findings from the Open Field Test demonstrated that repeated sub-anesthetic doses of ketamine induce significant hyperlocomotion in a mouse model of schizophrenia. The increased movement activity and elevated rearing events in the ketamine-treated mice suggest heightened motor activity, which is characteristic of ketamine’s psychostimulant properties [17, 18]. Cognitive function, as inferred from the EPM performance, also appeared to be impaired in the ketamine-treated mice. The ketamine group demonstrated prolonged latency to enter the open arms (p < 0.05), which may reflect deficits in decision-making processes associated with cognitive dysfunction in schizophrenia [14, 15]. Alteration in exploratory behaviour and preference for the closed arms in the ketamine-treated mice in elevated maze plus (EMP) is consistent with previous findings linking ketamine administration to altered cognitive function in schizophrenia models (13, 19].

The increase in SOD activity in these models is of particular interest as it may provide insights into the oxidative mechanisms underlying schizophrenia and the potential neuroprotective or neurotoxic roles of elevated SOD. In this study, we investigated the impact of repeated sub-anesthetic doses of ketamine on superoxide dismutase (SOD) levels in a mouse model of schizophrenia. Our results demonstrated that chronic administration of ketamine significantly elevated SOD activity, suggesting an enhanced oxidative stress response in schizophrenic mice. These findings align with existing literature that highlights the role of oxidative stress in schizophrenia pathology and the potential of ketamine to exacerbate this condition under certain dosing regimens [20]. The increase in SOD may reflect a compensatory mechanism to counteract the elevated Reactive Oxygen Species (ROS) levels induced by ketamine’s action on the brain which could contribute to the exacerbation of neurobiological and behavioural symptoms in schizophrenia. Atypical antipsychotic drug, risperidone administered to treatment group significantly (p< 0.05) decreased the elevated superoxide dismutase (SOD) level which is consistently associated with the amelioration of the behavioural symptoms compare to the KET group, in line with decreased SOD levels and diminished symptoms reported in risperidone treated schizophrenic patients in clinical finding [21, 22].

## Conclusion

The elevated SOD levels observed may reflect a compensatory mechanism against increased oxidative damage, which could contribute to the exacerbation of neurobiological and behavioural symptoms in schizophrenia.

